# Liquid co-substrates repower sewage microbiomes

**DOI:** 10.1101/261339

**Authors:** Justus Hardegen, Adriel Latorre-Pérez, Cristina Vilanova, Thomas Günther, Claudia Simeonov, Manuel Porcar, Olaf Luschnig, Christian Abendroth

## Abstract

A range of parameters are known to shape the methanogenic communities of biogas-producing digesters and to strongly influence the amount of biogas produced. In this work, liquid and solid fractions of grass biomass were used separately for semicontinuous batch methanation using sewage sludge as seed sludge. During 6 months of incubation, the amount of input COD was increased gradually, and the underlying methanogenic microbiome was assessed by means of microscopy-based automated cell counting and full-length 16S rRNA high-throughput sequencing. In this sense, we prove for the first time the suitability of the ONT™MinION platform as a monitoring tool for anaerobic digestion systems. According to our results, solid-fed batches were highly unstable at higher COD input concentrations, and kept *Methanosaeta* spp. typically associated to sewage sludge-as the majoritary methanogenic archaea. In contrast, liquid-fed batches developed a more stable microbiome, proved enriched in *Methanosarcina* spp, and resulted in higher methanogenic yield. This work demonstrates the high repowering potential of microbiomes from sewage sludge digesters, and highlight the effectiveness of liquefied substrates for increasing biogas productivity in anaerobic digestions.

## 1. Introduction

Anaerobic digestion is a well-known technology that allows microbial conversion of biomass into methane and carbon dioxide. Basically, anaerobic fermentation consists of four phases (Bischofsberger et al., 2005): hydrolysis (biomass fragmentation), acidogenesis (formation of organic acids, alcohols, carbon dioxide, and hydrogen), acetogenesis (formation of acetic acid), and methanogenesis (last phase of the process, in which acetic acid, hydrogen, and carbon dioxide are the main substrates for the formation of methane). The key microorganisms in the methanogenesis phase are methanogenic archaea, whose composition depends on the operation conditions and strongly changes when co-digestion with additional substrates occurs (Sundberg et al., 2013). Mesophilic methanogens that can be found in especially high abundances belong to the genus *Methanosaeta, Methanoculleus,* and *Methanosarcina* (Abendroth et al., 2015; Abendroth et al., 2017a). A strong microbial shift can be observed under thermophilic conditions, in which methanogens such as *Methanothermobacter* or *Methanobacterium* show an increased abundance (Maus et al., 2016; Lin et al., 2017; Xiao et al., 2018). Besides temperature, methanogens are also very sensitive to the organic loading rate. For example, under mesophilic conditions digestion processes with high amounts of chemical oxygen demand (COD) tend to have high amounts of *Methanosarcina* and *Methanoculleus.* On the other hand, sewage sludge, which has typically lower amounts of COD compared to typical industrial co-digesters, tends to have higher amounts of archaea corresponding to the genus *Methanosaeta* (Abendroth et al., 2015; Abendroth et al., 2017a).

Even though there is a basic understanding of the distribution of methanogenic genera under certain digestive conditions, there are still some gaps remaining. For example, to the best of our knowledge, the gradual increase of COD in sewage sludge by means of co-digestion with other substrates has not been sufficiently characterized. According to the current state of art, the mentioned microbial transition is of high interest for the scientific community in this field, as this process is of crucial importance for an efficient repowering of sewage digesters of municipal water treatment plants. With aims to reach the climate objectives of the European Union for the 21th century, researchers are continuously investigating new technologies and methodologies that might help to build a green and self-sustainable economy. In this context, a very promising approach is to increase the efficiency of water treatment plants by using existing sewage sludge digesters, in which co-digestion is implemented. Such a technological upgrade would allow efficient and local usage of organic waste sources, which are produced by surrounding communities and industries. In addition, it would be a further step towards powering self-sufficient water treatment plants. Based on this idea, a number of works showing the possibility to repower sewage sludge digesters by using cosubstrates have been published recently. The proposed co-substrates include grass biomass (Hidaka et al., 2016; Abendroth et al., 2017), food waste (Zahan et al., 2016), municipal solid waste (Cabbai et al., 2016), glycerol (Jensen et al., 2014), microalgae (Mahdy et al., 2015), or pear residues (Arhoun et al., 2013). To make such repowering approaches applicable for the industry, and to meet the high standards of water treatment plants regarding process stability, a better understanding of the microbial changes ocurring during the transition from typical sewage digestion to high-load digestion processes is still needed. This work aimed to investigate the impact of slowly increasing concentrations of COD on the underlying microbiome of sewage sludge digesters. The impact of different feeding strategies (feeding with liquids or solids) was also analysed. On the one hand, lignocellulose from fresh grass biomass was mechanically treated, separated from liquids, and used for the solid feeding strategy. On the other hand, grass liquor (after separation from the solid fraction), was used for the liquid feeding strategy.

A powerful tool to investigate microbiome changes is 16S rRNA gene amplicon high throughput sequencing or shotgun metagenomic approaches (e.g Vanwonterghem et al., 2014; Abendroth et al., 2017b), since they enable the detection of thousands of species in one single experiment, and can also yield information on the metabolic pathways underlying the biogas production process. Within the present work, we aimed to apply the recently developed ONT™MinION sequencing platform, as this technology could have the potential to become a suitable monitoring tool for anaerobic digestion plants. The use of metagenomic sequencing as a monitoring tool for industrial processes (i.e.: fermentations) has not been sufficiently explored to date. One of the main reasons for this is the economic investment needed to acquire a sequencer, as well as the technical complexity of the sequencing process and the ulterior bioinformatic analysis. This process is typically simplified by submitting samples to specialized sequencing facilities. Unfortunately, this makes the whole procedure significantly slower (results are typically obtained after some weeks). However, the launch of the ONT™MinION sequencer opens up a new scenario for real-life sequencing applications. Therefore, the features of this technology (user-friendly operation, real-time analysis, and portability) prove the unprecedented impact of ONT™MinION sequencing in the clinical (Quick et al., 2017), biosecurity (Pritchard et al., 2015), and environmental (Brown et al., 2017) fields. This work assesses for the first time the suitability of the ONT™MinION platform as a monitoring tool for anaerobic digestion systems, and uses this technology to follow up the changes in the archaeal methane-producing community at nearly real time.

## 2. Material and Methods

### 2.1 Chemical analysis and sampling

Fresh grass biomass was chosen to be used as substrate (Gramineae). Grass was pre-treated using a conventional juicer (Angel Juicer 8500 s, Angel Co.LTD., Corea). The solid fraction contained a COD of 366 mg O_2_ per g of substrate (according to the German guideline DIN 38414-S9), and the liquid fraction had a COD of 82,200 mg/L (according to the German guideline DIN 38409-H41). The amount of total volatile fatty acids (TVFA) and the solubilisation of COD were monitored every two or three days using conventional photometer-based assays (Nanocolor CSB15000 and Nanocolor organische Suren 3000, Macherey-Nagel, Germany). Organic loading rate was adjusted at the beginning of the experiment in such a way that the liquid fed reactors (B and C) produced a similar amount of methane as the solid fed reactors (D and E). Then, it was increased throughout the experiment, as shown in Figure 1. Produced gas was analysed using the “COMBIMASS GA-m” gas measurement device (Binder, Germany) to determine the ratio of CO_2_ and CH_4_.

**Figure 1:**
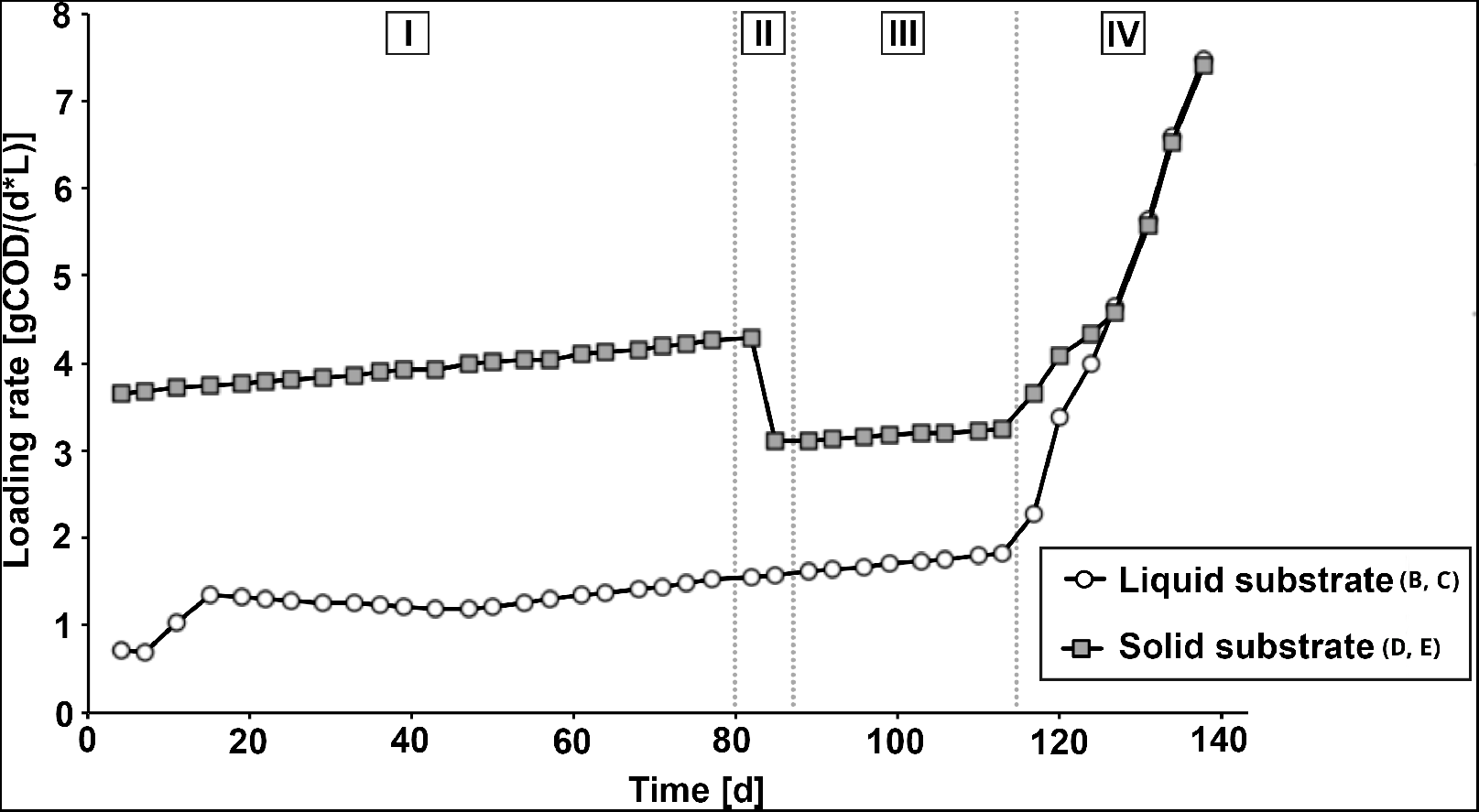
Substrate input in time. The addition of liquid (batch reactions B and C) and solid (batch reactions D and E) substrate was performed in four different phases, as indicated in roman numbers. Phases I and III: COD input concentrations were adjusted to a value in which similar amounts of biogas (methane) were produced in batches fed with liquid or solid substrates. Phase II: solid-fed batches (reactions D and E) reached an extremely high viscosity, and small amounts of water were added to enable stirring. Phase IV: in order to drastically increase the organic loading rate, the substrate was changed to molasses in all the reactors.

To compare the buffer capacity and the capacities for an increase of loading rates, the ratio of VFA and total inorganic carbon (TIC) was measured occasional using an automatic titration device following the instructions of the manufacturer (Biogas Titrator for FOS/TAC analyses, Hach-Lange, Germany).

### 2.2 Semicontinuous batch digestion

Five batch reactors were set up according to the German guideline “VDI 4630”. Reactor A was used as a negative control (without substrate input); reactors B and C were fed with liquid substrate (liquids separated from grass biomass); and reactors D and E were fed with solid substrate input (solids separated from grass biomass). Each reactor was filled with 300 mL of sewage sludge as seed sludge from the water treatment plant in Jena (Germany). The reactors were opened every 3-4 days - twice a week - to take samples and to add substrate. Afterwards, the reactors were closed and flushed with nitrogen to ensure an anaerobic atmosphere. Gas was collected in a liquid *7* displacement device (eudiometer) and measured daily.

In the beginning (day 1-11), the loading rate was adjusted in such a way 0 that liquid- and solid-fed reactors produced similar amounts of methane. After running the reactors with a constant input, at day 47 the organic loading rate for the liquid-fed reactors was increased by 0.4 mL per cycle. This was due to a ratio of volatile fatty acid to total inorganic carbon (VFA/TIC) of 0.178, which was much lower than the VFA/TIC in the solid-fed batches (0.441), indicating a higher capacity of substrate degradation in the liquid-fed batches.

From day 113 onwards, the input of both liquid and solid substrates was 9 reduced by 25 % each cycle, reaching a reduction of 100 % at day 124. At the same time, the grass biomass was replaced with molasses to induce a shock loading. The molasses input was increased by 0.5 g per cycle. Additionally, a volume of 100 mL of water was added to reactors D and E at day 82, as the sludge was too dense to be stirred.

### 2.3 Fluorescent microscopy

Prokaryotes were quantified after staining with 4’,6-diamidino-2-phenylindole (DAPI) using a epi-fluorescent microscope (Axio Lab.A1, Carls Zeiss, Germany). Firstly, teflon-coated slides with 10 wells (Carl Roth, Germany) were 3 covered with a gelatine membrane. To do this, 10 *μ*L of gelatine solution (0.1 9 *%* gelatine und 0.01 *%* CrK(SO4)2) was pipetted on each well and dried at 50°C for 10 minutes. Depending on the density of cells, samples were diluted 1:200 or 1:2000 with PBS buffer. Each well was filled with 10 *μ*L of the diluted sample and dried at 50 °C for 10 minutes. Finally, 2.5 *μ*L of fluorescent mounting solution (Roti®-Mount Aqua, Carl-Roth, Germany) was applied. Quantification was performed under 400x magnification and 450 ms exposure time. For each time point, 32 pictures ware taken and evaluated using the ImageJ software.

Methanogenic archaea were quantified using the same microscope, but with a different set of optical filters and an excitation wavelength adjusted to the quantification of the cofactor F420 (which is associated with methanogenic archaea). Samples were diluted 1:2 with a mounting solution (10 *μ*L each) (Roti®-Mount FluorCare, Carl-Roth, Germany) and 3 *μ*L of the suspension was applied between the cover slip and the slide. Due to increasing concentrations of total solids in the batch bottles D and E, a 1:10 pre-dilution was applied for these samples after several weeks. Pictures were taken with 400x magnification and 126 ms exposure time. For each time point, 48 pictures ware taken and evaluated using the ImageJ software.

### 2.4 DNA isolation

In order to reduce the amount of inhibiting substances, biomass samples were sedimented by centrifugation (510 min at 20,000 g) and washed several times with sterile PBS buffer until a clear supernatant was observed. Then, metagenomic DNA was isolated using the Power Soil DNA Isolation kit (MO BIO Laboratories) following the manufacturers instructions. The quantity and quality of the DNA was determined on a 1.5 *%* agarose gel and with a Nanodrop-1000 Spectrophotometer (Thermo Scientific, Wilmington, DE).

### 2.5 16S rRNA gene amplification and barcoding

The full-length 16S rRNA gene of archaea was PCR-amplified using universal primers Arch8F (5’-TCCGGTTGATCCTGCC-3’) and Arch1492R (5’‐ GGCTACCTTGTTACGACTT-3’), for which specificity had been previously reported (Klindworth et al., 2013). Primer sequences were tailored to add the ONT™Universal Tags (5’-TTTCTGTTGGTGCTGATATTGC-3’ for the forward primer and 5’-ACTTGCCTGTCGCTCTATCTTC-3’ for the reverse primer) to their 5’ ends. These universal tags allowed the barcoding of the amplicons in the second PCR using the ONT™PCR Barcoding kit (EXPPBC001).

For the first PCR, the mixture consisted of 1X Taq Polymerase Buffer, 200*μ*M dNTPs, 200nM primers, 1 U of Taq DNA polymerase (VWR), and 10ng of DNA template in a final volume of 50*μ*L. PCR conditions were an initial denaturation step at 94°C for 1min, followed by 35 cycles of amplification (denaturing, 1min at 95°C; annealing, 1min at 49°C; extension, 2min at 72°C), with a final extension at 72°C for 10min. To assess possible reagent contamination, each PCR reaction included a negative control without template DNA, which did not amplify. A purification step using Agencourt AMPure XP beads (Beckman Coulter) at 0.5X concentration was performed to remove primer-dimers and non-specific amplicons, and the resulting DNA was recovered and assessed by Qubit quantification.

In the second PCR, the mixture contained 0.5 nM of the first PCR product, 1X Taq Polymerase Buffer, 200M of dNTPs, 1 U of Taq DNA polymerase (VWR), and the corresponding specific barcode (EXP-PBC001) as recommended in the ONT protocol 1D PCR barcoding amplicons (SQK-LSK108). The PCR conditions consisted of an initial denaturation step for 30 s at 98 C, followed by 15 cycles at 98°C for 15s, 15s at 62 °C for annealing, 45s at 72°C for extension, and a final extension step for 7min at 72°C. A clean-up step using AMPure XP beads at 0.5X concentration was used again to discard short fragments as recommended by the manufacturer. Finally, an equimolar pool of amplicons was prepared for the subsequent library construction.

### 2.6 Library construction and sequencing

The Ligation Sequencing Kit 1D (SQK-LSK108) was used to prepare the amplicon library to load into the MinION™following the instructions of the 1D PCR barcoding amplicon protocol of ONT. The barcoded pool of amplicons (1 μg) was used as input DNA. The DNA was processed for end repair and dA-tailing using the NEBNext End Repair / dA-tailing Module (New England Biolabs), and the resulting DNA was purified using Agencourt AMPure XP beads (Beckman Coulter) and assessed by Qubit quantification. For the adapter ligation step, a total of 0.2 pmol of the end-prepped DNA was added to a mix containing 50 *μ*L of Blunt/TA ligase master mix (New England Biolabs) and 20 *μ*L of Adapter Mix (SQK-LSK108), and was incubated at room temperature for 10min. DNA was purified again with the Agencourt AMPure XP beads (Beckman Coulter) and the Adapter Bead Binding buffer provided on SQK-LSK108 kit to finally obtain the DNA library.

The flow cell (R9.4, FLO-MIN106) was primed and then loaded as indicated in the ONT™protocols. Sequencing was performed during 12h using the standard sequencing protocol implemented in the MinKNOW™software.

### 2.7 Metagenomic data analysis

Reads were basecalled using the Metrichor™agent, and sequencing statistics were followed in real time using the EPI2ME debarcoding workflow. The fast5 files obtained were converted to fastq files using poRe (Watson et al., 2015) and adapters were trimmed using Porechop (https://github.com/rrwick/Porechop). The resulting sequences were analyzed with the QIIME software. Briefly, reads were aligned, and then identified through BLAST searches against the latest version of the GreenGenes database (13_8). Data was then further analyzed and represented with custom scripts. In order to detect significant changes in the relative abundance of particular taxa *(Methanosaeta* spp. and *Methanosarcina* spp.), a Welchs t-test for unequal variances was performed.

## 3 Results and Discussion

### 3.1 Reactor performance: liquid vs. solid feeding

Two different strategies for the repowering of sewage sludge involving co-digestion were compared: (1) using a liquid co-substrate with very low percentage of total solids (TS) and, therefore, with low amounts of ligno-cellulose (batch reactions B and C); and (2) using a solid co-substrate with a very high percentage of TS and, therefore, with high amounts of ligno‐ cellulose (batch reactions D and E). Both co-substrates were obtained from fresh grass (Graminidae) biomass. Additionally, a control digester was kept without co-substrate input (batch reaction A). The loading rate of both repowering approaches was adjusted in such a way that both systems produced similar volumes of methane per working volume and that the concentration of solubilized COD and TVFA was increasing slowly, as described in Material and Methods (Fig. 1 and 2).

**Figure 2:**
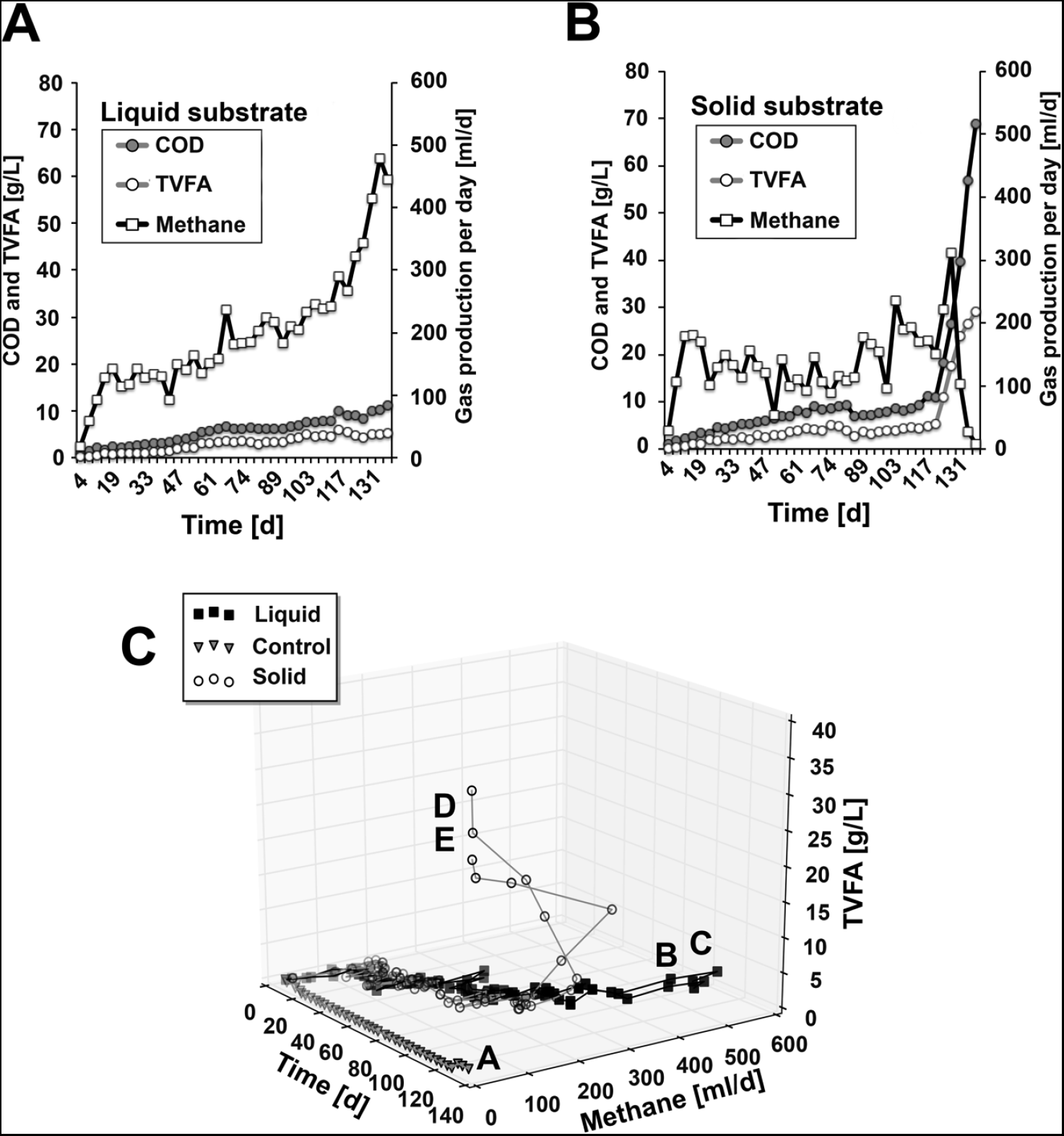
Chemical parameters measured for the different experimental set-ups. Data on COD, TVFA and methane production are represented for one of the liquid fed reactors (**A**) and one of the solid fed reactors (**B**). A tridimensional representation of the evolution of all the reactors is also shown (**C**).

In both repowering strategies, the amount of biogas ranged between approximately 100 and 200 mL of methane per day during phase I (day 1 – 82) (Fig. 2). By the end of phase I the liquid fed batches reached a solubilized COD of 5.9 ± 0.2 gCOD/L, and a TVFA concentration of 2.81 ± 0 gTVFA/L. Although the solubilized COD and TVFA of the solid fed system were higher at that time (12.02 ± 2.98 gCOD/L, and 3.98 ± 0.02 gTVFA/L), the produced amount of methane was slightly higher in the liquid fed system (Fig. 2A and B). Moreover, methane production within the liquid fed system proved more stable in time. By the end of phase I, the digestion sludge in the solid fed system reached such high viscosity that no stirring was possible. In order to ensure a better substrate distribution and to facilitate to movement of bubbles, a small amount of water was added to the solid fed batch systems D and E (100 mL). Due to the dilution, the loading rate of the solid fed batches was slightly reduced during phase 3 (Fig. 1). Altogether, the described observations indicate that the liquid fraction was easier to dose and pump, more predictable, and more stable (also, solid layers of scum were not formed on the surface).

Since the high viscosity of the solid fed batch prevented any further increase of the loading rate, the substrate was changed stepwise to molasses for both repowering experiments (liquid and solid fed batches), starting at day 117. In parallel, the loading rate of the other substrates (liquid and solid grass biomass) was lowered stepwise until day 127, when both experimental set-ups were fed exclusively with molasses.

The solubilized COD increased drastically in both repowering approaches, indicating a substrate overload. However, from day 132 onwards (Fig. 1, phase 4), the produced amount of methane became drastically reduced in the solid fed batches (Fig. 2B and C). In contrast, the liquid fed batch systems displayed higher stability, with continuously increasing levels of methane production. Moreover, a sudden acid shock was detected in the concentration of TVFAs in the solid fed batches, reaching more than 20 gTVFA/L at day 132 (Fig. 2B). The liquid fed batches remained with a lower concentration of TVFAs, reaching 5.52 ± 0.54 gTVFA/L at day 32, indicating a much more efficient conversion of TVFAs into methane.

### 3.2 Changes in the abundance of prokaryotes and methanogenic archaea

Fluorescent microscopy was performed to complement the chemical analysis during the experiments. As described in Material and Methods, DAPI staining was used to count the total number of prokaryotes, and fluorescence associated to the cofactor F420 was used to quantify methanogenic archaea (Fig. 3). The number of prokaryotes varied between 1E+09 and 1E+10 per mL, which is comparable with previous studies (Nettmann et al., 2010). The feeding events in both liquid‐ and solid-fed batches did not cause a noticeable increase in the number of prokaryotes. The substrate overload at the end of the experiment (Fig. 1, phase 4) did not cause a shift in the number of prokaryotes, indicating a high stability of the underlying bacterial community. However, starvation in the unfed control (batch reaction A) caused a decrease in the number of prokaryotes below 1E+09. A similar influence of starvation was observed recently in another study (Abendroth et al., 2016).

**Figure 3:**
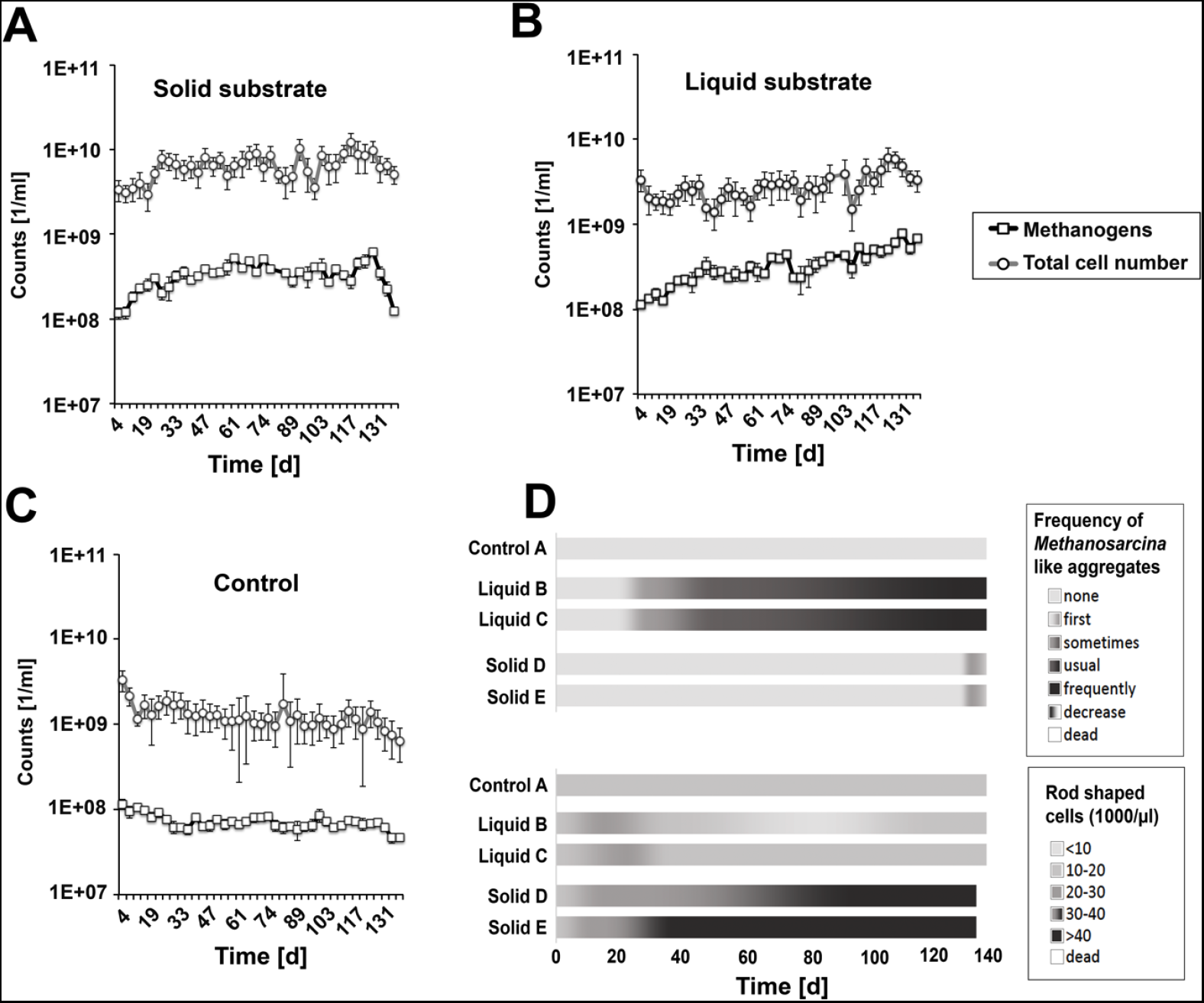
Microscopic analysis of prokaryotic and methanogenic communities. Methanogens were screened by quantifying the co-factor F420, whereas total Prokary-ota were stained with DAPI. Quantified F420‐ and DAPI signals are shown for a liquid-fed reactor (**A**), a solid-fed reactor (**B**), and the unfed control (**C**). Additionally, *Methanosarcina* spp. like cell aggregates and rod shaped F420-signals were analysed semiquantitatively (**D**). High amounts of rod shaped F420-signals were used as indicators for high concentrations of *Methanosaeta*, which is typical for sewage plants and sludges with low COD content.

The number of methanogens increased continuously from 1E+08 to 1E+09 per mL, which is in accordance with other studies (Nettmann et al., 2010). However, in the unfed control the number of methanogens remained unexpectedly stable, indicating a high resistance against starvation. At day 132, the number of methanogens decreased back to 1E+08 cells in the solid-fed batches, but remained stable in the liquid-fed batches. This is in accordance with the halt in methane production observed in the solid fed batches at day 132 (Fig. 2). Taken together, these results suggest the presence of a better-adjusted microbiome in the liquid fed batches.

### 3.3 Changes in the composition of the methanogenic microbiome

Changes in the relative abundance of the main genera involved in methane production were followed up by means of microscopy and confirmed by full length 16S rRNA high-throughput sequencing, using specific primers targeting archaea. As shown in Fig. 3D, microscopic analysis revealed a high number of rod-shaped methanogens in solid-fed batches, indicating a high number of *Methanosaeta,* a typical genus observed in sewage sludge (Abendroth et al., 2015; Abendroth et al., 2017a). Interestingly, liquid-fed batches showed decreasing numbers of rod-shaped methanogens and increasing numbers of *Methanosarcina*-like complexes. Microscopic analysis made it difficult to quantify *Methanosarcina* species, as they tend to form large complexes which prevent the identification of single cells.

In parallel, the archaeal communities present in the different reactor configurations were analysed by means of full-length archaeal 16S rRNA high-throughput analysis using ONT^™^MinION sequencing. In accordance with microscopic data, the principal component analysis performed with the archaeal profile of each sample showed a different microbial evolution for solid‐ and liquid-fed reactors (Fig 4B). Figure 4A shows the evolution of the taxonomic composition in each experimental condition. At day 29, *Methanosaeta* was the most abundant genus in all cases, accounting for 23-75% of sequences. *Methanosaeta* remained the majoritary genus throughout the experiment in solid-fed batches, as well as in the control digester. However, the abundance of *Methanosarcina* spp. increased in liquid-fed batches, with it becoming the majoritary genus at days 103 (batch C) and 139 (batch B and C). Figures 4B and 4C show the enrichment of *Methanosarcina* spp. in liquid-fed reactors with respect to solid-fed reactors, and the opposite trend, observed for *Methanosaeta* spp.

**Figure 4:**
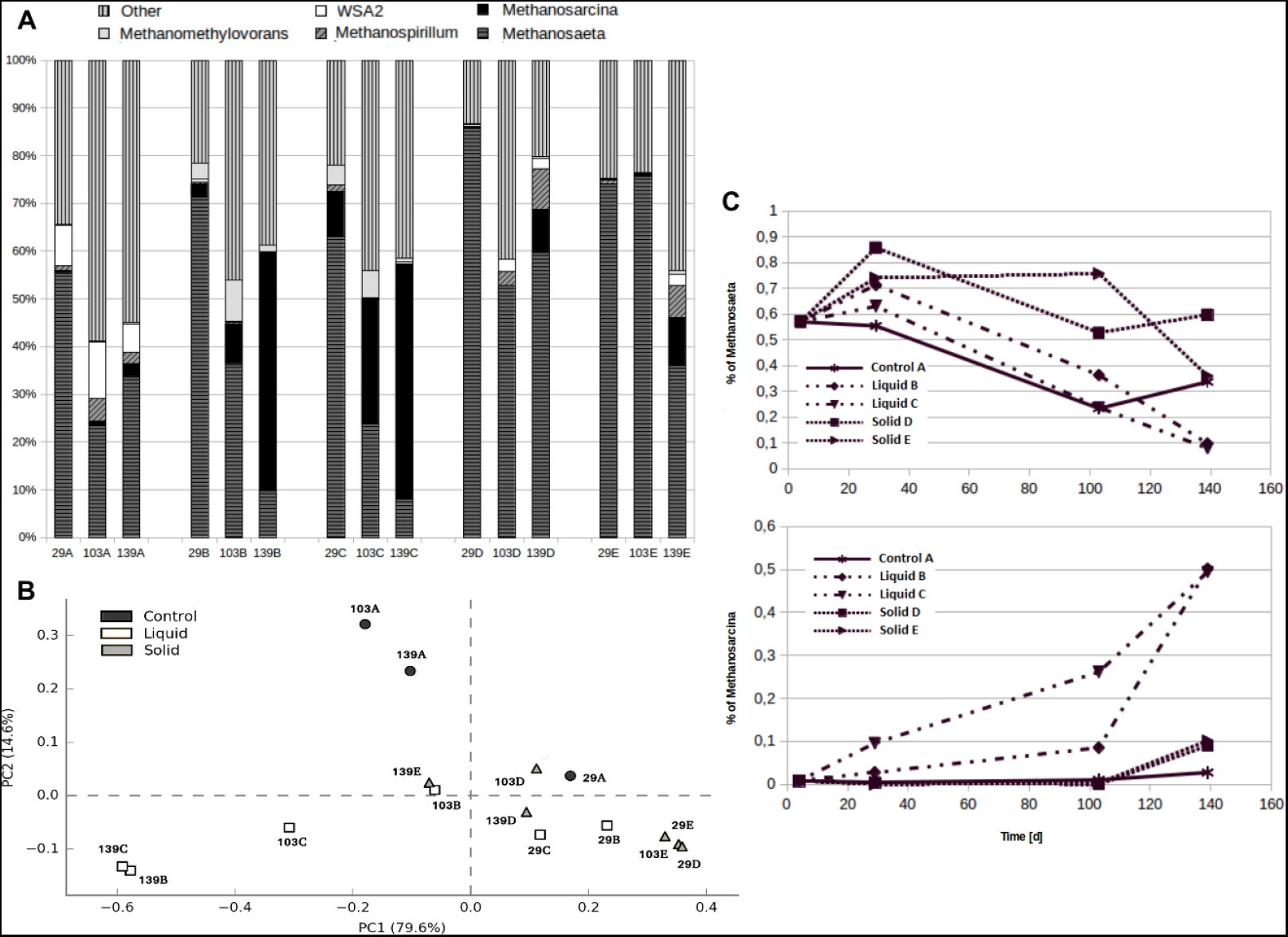
Taxonomic analysis of the archaeal communities based on full-length 16S rRNA massive sequencing. (**A**) Relative frequency of the most abundant taxa in each sample. Numbers in the X axis indicate the day in which samples were taken. Letters indicate the type of feeding (A: unfed control; B and C: liquid-fed batches; D and E: solid-fed batches). (**B**) Principal Component Analysis (PCA) based on the taxonomic profile of the samples. (**C**) Evolution of the frequency of *Methanosaeta* spp. (up) and *Methanosarcina* spp. (down) in the different reactors.

It has to be noted that *Methanosarcina* spp. are methanogens which can be found in co-digesters with high loading rates, especially in the leachate of leach-bed systems (Klocke et al., 2008; Zhao et al., 2013; Abendroth et al., 2015). Therefore, the observed results indicate that the methanogenic microbiome of liquid-fed systems can be successfully shaped and adapted to higher loading rates. This is in contrast with solid-fed systems, which remained enriched in *Methanosaeta* spp., a typical genera from sewage sludge digesters, which tend to have lower loading rates (Abendroth et al., 2015).

## 4. Conclusions

With the aim of investigating different strategies to use co-substrates in sewage sludge digesters, the impact of liquid and solid substrates on the underlying microbiome was investigated. To follow up changes in the archaeal communities associated with the different reactor configurations, chemical analysis, microscopic fluorescence-based quantification, and 16S rRNA high-throughput sequencing were applied to samples taken throughout the experiment.

For the first time in this field, 16S rRNA high-throughput sequencing was performed with an ONT™MinION sequencer. This technology enabled the analysis of full-length (1.5 kb) reads of the 16S rRNA gene from the archaeal communities, and most importantly, allowed us to obtain taxonomic data at nearly real time. Hence, ONT™MinION sequencing can be seen as a promising technology to be applied in real-world problems associated to the biogas industry, such as the real-time analysis of changing methanogenic communities, which play a central role in the efficiency of the production process.

According to our results, solid-fed batches proved unstable in terms of TVFAs production and conversion to methane, and were also difficult to dose and pump. Instead, liquid-fed batches displayed a more predictable and stable behaviour. Most importantly, the microbiome of liquid-fed reactors was re-shaped, changing from a *Methanosaeta* spp.-dominated community, to a *Methanosarcina* spp.-enriched community. In contrast, solid-fed batches remained dominated by *Methanosaeta* spp., which are typical for digesters with low organic loading rates.

Altogether, our results indicate that the addition of liquid co-substrates results in a more effective repowering of the methanogenic microbiome of sewage sludge, and results in higher biogas production. This paves the way towards a more efficient configuration of water treatment plants.

## Conflict of interest

The authors declare no conflict of interest.

